# A novel FRET-force biosensor for nucleoporin gp210 reveals that the nuclear pore complex experiences mechanical tension

**DOI:** 10.1101/2025.05.14.654032

**Authors:** Peipei Wang, Kevin Denis, GW Gant Luxton, Daniel E. Conway

## Abstract

The nuclear pore complex (NPC) is a large multi-protein structure that enables movement of macromolecules, such as mRNA and proteins, between the nucleoplasm and cytoplasm. There has been great interest in how the physical state of the NPC can influence nuclear-cytoplasmic transport. The hypothesis that the NPC may be mechanosensitive is supported by prior reports showing that the diameter of the NPC increases with nuclear envelope stretch as well as increased ECM stiffness. We therefore sought to develop a biosensor-based approach to determine if the NPC experiences mechanical tension. Using a previously developed FRET-force biosensor, known as TSmod, we developed a gp210 tension sensor. gp210 is a transmembrane nucleoporin, which may serve to anchor the NPC into the nuclear envelope. Using a CRISPR knock-in strategy, we developed a HeLa cell line which expresses the gp210 tension sensor at endogenous levels. Using this sensor, we observed that the biosensor exhibited FRET changes that were consistent with increased force in response to osmotically induced nuclear swelling. Cell attachment, the nuclear LINC complex, ECM stiffness, chromatin condensation, and actomyosin contractility were all observed to influence gp210 forces. Surprisingly, gp210 forces were increased with chromatin relaxation and myosin light chain kinase inhibition, indicating that NPC forces may be differentially regulated from forces on the LINC complex. Our data support a hypothesis where nuclear strain, rather than cytoskeletal forces, is the predominant source for NPC forces. Our studies provide the first direct measurements of protein-level mechanical forces on the NPC. We anticipate that the gp210 force sensor will be of use for future studies of NPC mechanobiology.

## Introduction

The nuclear pore complex (NPC) serves as the major gateway for macromolecules (mRNA and proteins) moving between the cytoplasm and the nucleus. The NPC is formed by approximately 30 different proteins, known as nucleoporins (nups). These nucleoporins are organized into different regions of the NPC, including the central channel that is formed by phenylalanine- and glycine-rich nucleoporins, known as FG-nups. Macromolecules moving from the cytosol to the nucleus (or vice versa) must pass through this central channel. While small molecules less than 5-9 nm in diameter can pass through the NPC, larger molecules require interactions with nuclear transport receptors (importins and exportins) to move through the NPC. The ability of importins and exportins to shuttle these macromolecules through the NPC is further dependent on Ran GTPase. Classically, regulation of transport of these macromolecules is thought to occur by chemical modifications to the macromolecule (e.g. mRNA polyA tail, protein phosphorylation) that affect their affinity for nuclear transport receptors. Thus, the NPC was not initially considered to be a regulator of the kinetics of nuclear-cytosolic transport beyond serving as a barrier to unrestricted movement.

Changes in the size of the central channel of the NPC have the potential to affect the rate of nuclear-cytosolic transport. The NPC central channel exists in two states: constricted (40-50 nm) and dilated (55-70 nm).^1^ The hypothesis that mechanical forces could modulate the size of the central channel was proposed by both Mohammad Mofrad^2^ and Don Ingber.^3^ The first experimental evidence to support this hypothesis came from work by the Roca-Cusachs group which showed that the size of the NPC increases when cells are cultured on increasing ECM stiffness.^4^ Furthermore, they showed that mechanical forces can control both passive and facilitated nucleocytoplasmic transport.^4,5^ An additional study by the Beck group showed that the mechanical state of the nuclear envelope controls NPC diameter.^6^

Despite the prior work showing a positive correlation between force and NPC size, direct evidence that NPC proteins (e.g. nucleoporins) experience mechanical tension is lacking. In this study we developed a FRET-based tension sensor for gp210 (nup210), a transmembrane NPC protein, using a previously developed FRET-force module known as TSmod.^7^ Using CRISPR-cas9, we developed a knock-in gp210 tension sensor cell line so that the sensor would be expressed at endogenous levels. Using this cell line, we demonstrate that gp210 experiences significant mechanical tension. Using hypo- and hyper-osmotic shock experiments we show the gp210 tension is dependent on nuclear envelope tension. Additional factors which regulate gp210 forces include attachment to ECM, ECM stiffness, DNA condensation, and myosin contractility. Some perturbations such as chromatin relaxation (Trichostatin A) or myosin inhibition (ML-7) resulted in opposite changes on gp210 forces as compared to prior studies of nesprin ^8^ and lamin A/C forces,^9^ which suggest that gp210 forces may also be influenced by changes in nuclear strain, as well as forces across the LINC complex. Finally, we showed that conditions which increase gp210 forces (hypo-osmotic shock) increase the rate of nuclear export. Taken together our findings directly demonstrate that NPC proteins do experience mechanical tension and that these forces can be correlated to changes in nuclear-cytosolic transport kinetics.

## Methods

### Cell culture

HeLa cells were obtained from ATCC (CCL-2) and cultured in DMEM with 10% FBS. For all imaging experiments, cells were grown on glass bottom dishes and allowed to adhere for one day.

### gp210 DNA construct

Rat Nup210-GFP was obtained from Martin Hetzer.^10^ Using standard cloning approaches the GFP was removed and replaced with TSmod^7^ to create gp210 TS. A force-insensitive control sensor, termed gp210 F7, was created by replacing the flexible 40 amino acid linker in TSmod with a flexible 7 amino-acid long peptide (F7) that was previously described.^11^ A second control sensor was made in which the transmembrane domain of gp210 was replaced with that of Sec61β, which was previously reported to inhibit the ability of gp210 to associate with nuclear pores.^10^

Cells were transiently transfected with these constructs using Lipofectamine 3000 Transfection Reagent (ThermoFisher Scientific).

### HeLa knock-in strategy

In collaboration with the Ohio State University Gene Editing Shared Resource, the generation of NUP210 TSmod Knock in HeLa cells was performed as previously described.^12,13^ Briefly, a gRNA designed against the human NUP210 gene (5’-ACTCACAAGGACCGGGCCCA-3’ cutting the gene coding sequence between aa 1791 and 1792) was cloned in the pX330-U6-Chimeric_BB-CBh-hSpCas9-hGem (1/110) plasmid (Addgene plasmid 71707).^14^ Then, we generated a donor plasmid containing the genomic sequence upstream (863 nt) and downstream (854 nt) of the gRNA cut site, flanking the TSmod sensor.^7^

HeLa cells were transfected with plasmids using Lipofectamine™ 3000 Transfection Reagent. After 48hours transfection, single cells were sorted into individual 96-wells plate to generate single cell clones using Aria III at the Analytical Cytometry Shared Resource of the Ohio State University Comprehensive Cancer Center. After 10-14 days, individual colonies were passaged and screened by PCR analysis. PCR reactions were carried out with 100 ng genomic DNA in the GoTaq Master Mix (Promega) according to the manufacturer’s instruction (Forward primer: 5’ TGCTTCCAGAGATGGCCATGCA 3’, Reverse primer: 5’ AAAACCACCAGGCCCACTGT 3’). A ∼2300bp amplicon was observed for the knock-in and a ∼700 bp amplicon for the wild-type sequence. Additional screening was performed by fluorescence microscopy (to confirm localization to the nuclear envelope) and fluorescence recovery after photobleaching (FRAP) (to confirm FRET).

### FRET image acquisition and analysis

Fluorescence lifetime imaging was performed using Leica STELLARIS FALCON confocal microscope equipped with Plan Apochromat ×40 water immersion objective. Cells were imaged on glass bottom dishes (#1.5 coverslip). Cells were excited with White Light Laser Stellaris 8 at 450□nm, and fluorescence lifetime times were recorded with HyD X detector, in the range 465 to 490□nm, 20% laser intensity, 7 frame repetitions to obtain the photon arrival times specific to donor emission. The pixel-by-pixel photon arrival times were fitted for bi-exponential decay components using n-Exponential Reconvolution fitting model of Leica LAS X software to obtain mean lifetimes from individual cells. Efficiency (E) was calculated using the following equation: E=1-*τ* /mean *τ* mTfp.

### Acceptor photobleaching experiments

Acceptor photobleaching experiments were performed using Leica STELLARIS FALCON confocal microscope equipped with Plan Apochromat ×40 water immersion objective. Cells were imaged on glass bottom dishes (#1.5 coverslip). Cells were excited with White Light Laser Stellaris 8 at 515□nm (100% laser intensity, 10 frame repetitions), and fluorescence measured using HyDS detectors (donor 460-590nm; acceptor 525-600nm).

### Immunostaining

NPCs were labeled using Mab414 (biolegend, 902907). Lamin A was labeled with c-terminal lamin A antibody (sigma L1293). Vimentin was labeled with vimentin E-5 antibody (Santa Cruz sc-373717). Actin was labeled with Acti-stain 670 phalloidin (Cytoskeleton). Cells cultured on 35mm glass bottom dishes were washed with PBS, then fixed with 4% paraformaldehyde 10min, wash with PBS 10min x 3, incubate in 5%BSA+0.15%TX100 PBS for 1hour, incubated with primary antibodies(1:200, 4degree overnight), wash with PBS, secondary antibodies(Alexa Fluor 568 donkey anti – rabbit or mouse IgG 1:400 RT 1 hour), wash with PBS, mounting with ProLong™ Gold Antifade Mountant with DNA Stain DAPI (ThermoFisher P36931).

### Osmotic stress experimental details

Hypo-osmotic shock experiments were performed as previously described.^4^ Briefly, cells were changed from normal (isotonic) cell culture media to hypotonic media (2/3 de-ionized water + 1/3 imaging medium). FLIM measurements were performed for 5-30min exposure to hypotonic media. For recovery experiments cells were transitioned back to isotonic media for the timepoints indicated.

Hyper-osmatic shock experiments were performed as previously described.^9^ Briefly, cells were changed from normal (isotonic) cell culture media to media containing 250mM sucrose. FLIM measurements were performed for 5-30min exposure to hypertonic media. For recovery experiments cells were transitioned back to isotonic media for the timepoints indicated.

### Small molecule inhibitors

The following small molecule inhibitors were used as indicated: Methylstat (Sigma, 2.5 μM 48h), Trichostatin A (TSA) (Sigma, 100nM, 24h), ML 7 (Cayman, 5 μM 30min), Y27632 (Cayman, 10μM 1.5h), Rho Activator II (Cytoskeleton, 0.25ug/ml 1h).

### Disruption of the LINC complex

To disrupt the LINC complex siRNA against SUN1 (UUACCAGGUGCCUUCGAAA) and SUN2 (UGGCAGAGAUGCAGGGCAA) was synthesized by Sigma. These sequences were previously shown to be effective for knockdown of SUN1 and SUN2.^16^ Cells were transfected with Lipofectamine™ RNAiMAX Transfection Reagent (ThermoFisher). Control cells were transfected with MISSION® siRNA Universal Negative Control #1 (Sigma, SIC001-1NMOL). Knockdown of SUN1 and SUN2 was validated using western blotting with anti-SUN1 (Abcam AB124770) and anti-SUN2 (Abcam AB124916).

### ECM experiments

In the indicated experiments to assess the role of the ECM, cells were seeded on uncoated glass bottom dishes (control), coated with 0.01% poly-l-lysine hydrobromide (Sigma, P4832), or with 20 µg/mL bovine fibronectin (Sigma F1141). In other indicated experiments cells were seeded on CytoSoft Imaging 24 well plates (Advanced Biomatrix) with stiffness of 2, 8 or 64 kPA.

### Nuclear morphology measurements

Nuclear cross-sectional area, and the major and minor axis length, for each nucleus (stained by Hoechst) was measured using ImageJ. Nuclear elongation was measured by the ratio of major axis length to minor axis length.

### Nuclear import experiments

NLS-mcherry-LEXY (Addgene plasmid 72655) was used to measure import kinetics as previously described.^15^ Cells were transfected with this plasmid using Lipofectamine 3000, and selected with G418 for stable expression. Cells expressing the LEXY biosensor were exposed to 488-nm laser power (30% 20s-50cycles) to induce nuclear export. Cells were then imaged using 568-nm (3% laser power). The intensity of the mcherry LEXY fluorescence was normalized to the intensity prior to photoactivation.

### Statistical Analysis

For statistical comparisons between groups of three or more conditions, one way ANOVA was used with the Geisser–Greenhouse correction, followed by post-hoc testing using Tukey’s multiple comparisons test with individual variances computed for each comparison. T-test was used for comparisons between two conditions. In all cases a value of p<0.05 was considered significant.

## Results

### Design of a gp210 tension sensor to measure NPC forces

We hypothesized that transmembrane nucleoporins would be most likely to experience mechanical tension. Previously, a gp210-GFP fusion protein was designed where the GFP was located at an internal insertion site.^17^ This gp210-GFP fusion protein has been shown to rescue cells in which endogenous gp210 was depleted,^10^ suggesting that insertion of fluorescent proteins at this site does not significantly alter gp210 functions. Using this strategy, we developed a gp210 tension sensor (gp210TS) in which the GFP was replaced with the FRET-force tension sensor TSmod^7^ (**Figure 1A**). HeLa cells transfected with gp210TS showed that the sensor localized to the nuclear envelope (**Supplemental Figure 1)**. We also observed increased donor fluorescence after acceptor photobleaching, indicating that this sensor experiences FRET (**Supplemental Figure 1)**.

**Figure 1.**
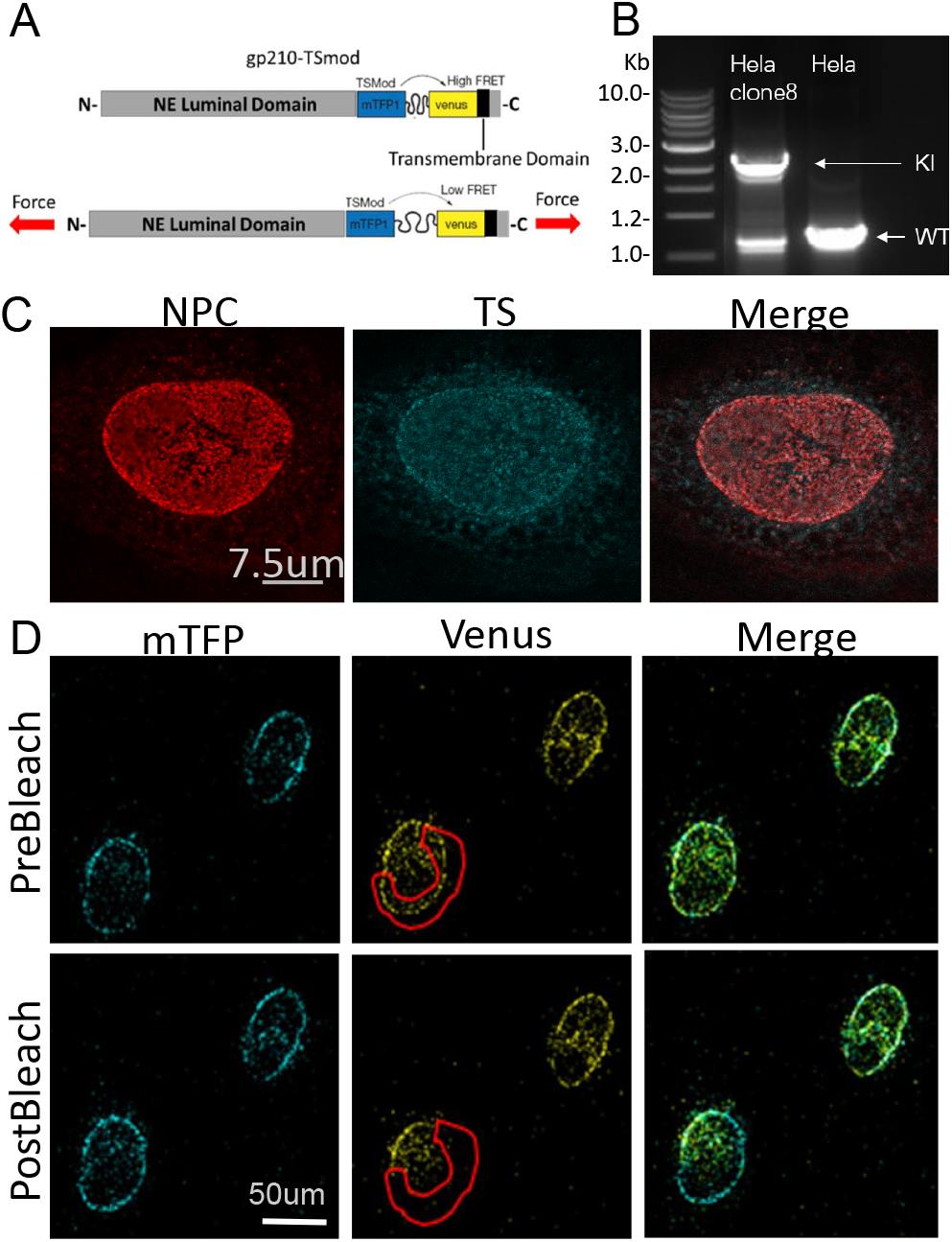
Design of a gp210 tension sensor to measure mechanical forces. **A)** Schematic of the gp210 tension sensor, in which TSmod was inserted at an existing internal site previously used for GFP-gp210. **B)** A CRISPR/cas9 strategy was used to knock in TSmod to the endogenous locus of gp210 in HeLa cells, where knock-in clones were identified as a larger PCR product. **C)** Immunostaining experiments showed that gp210TS was observed to co-localize with NPCs (NPCs were labeled using the pan-NPC mab414 antibody). D) An increase in donor (mTFP1) fluorescence was observed when the acceptor (venus) was photobleached, demonstrating FRET for the knock-in gp210 tension sensor.

Because overexpression of individual nucleoporins can significantly alter cellular phenotypes, we sought to develop a cell line where gp210TS was under the control of the endogenous promoter. Using CRISPR-cas9 to knock-in gp210TS, we developed a gp210TS HeLa cell line (**Figure 1B**). gp210TS was observed to co-localize with NPC immunolabeling (**Figure 1C**), indicating that gp210TS is incorporated into the NPC. Acceptor (venus) photobleaching resulted in increased donor (teal) fluorescence, demonstrating FRET is occurring in the gp210TS knock-in cell line **(Figure 1D**).

### Osmotic-induced changes in nuclear envelope tension affect gp210 forces

To test the responsiveness of the gp210 force sensor, we used hypo- and hyper-osmotic treatments to increase or decrease, respectively, nuclear envelope tension. Treatment of cells with hypo-osmotic shock increased nuclear area (**Figure 2A, 2C**) and decreased gp210 TS FRET efficiency (**Figure 2B**). We observed similar results with hypo-osmotic shock in transiently transfected HeLa cells (**Supplemental Figure 2**). The responsiveness of the gp210TS to osmotic swelling indicates that the gp210 tension sensor is responsive to changes in mechanical forces.

**Figure 2.**
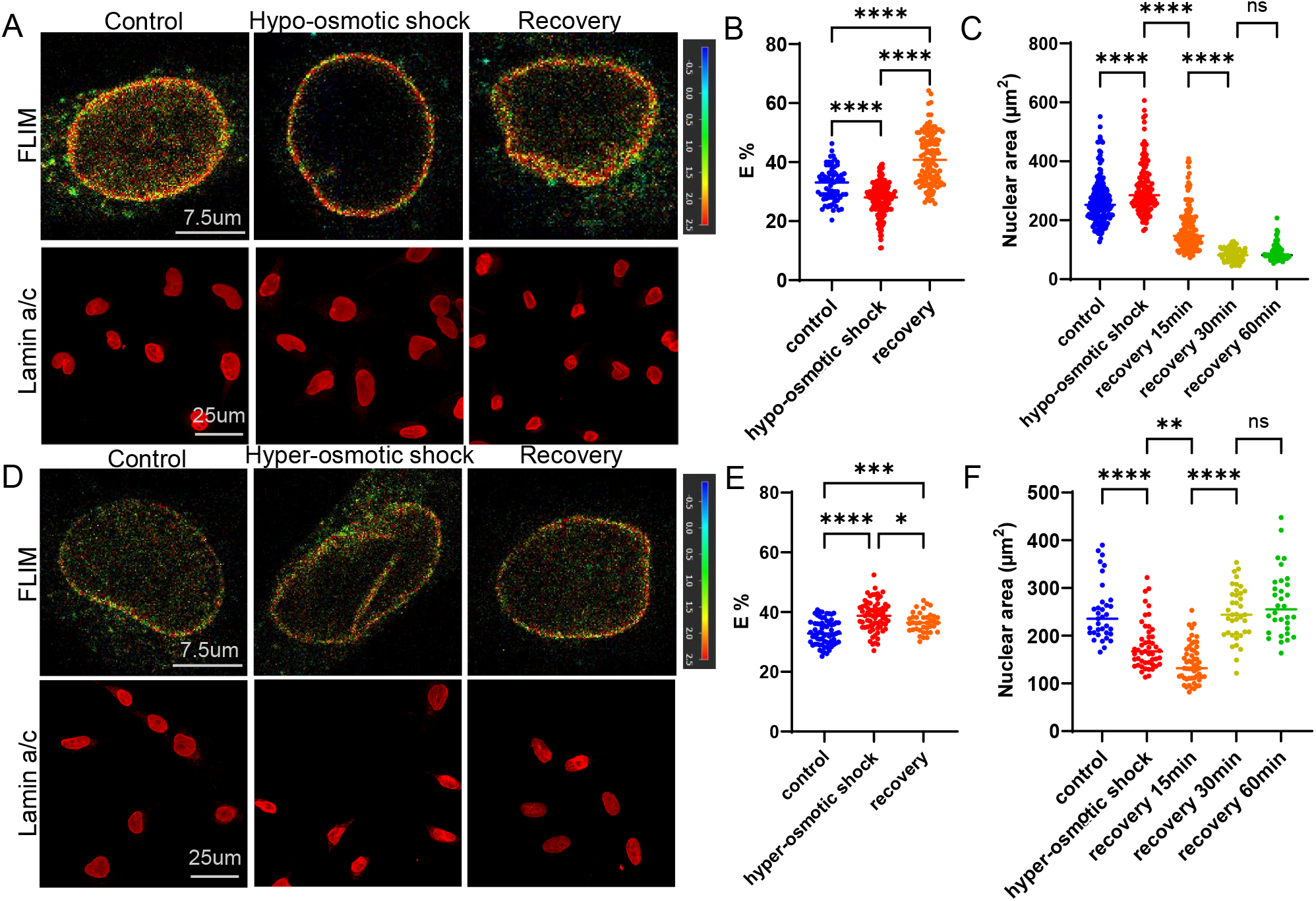
gp210 forces change during osmotic-induced changes in nuclear envelope tension. **A)** HeLa cells with gp210 tension sensor knock-in were exposed to isotonic (control) or hypo-osmotic buffer for 30 minutes to induce nuclear swelling. Additionally, a recovery sample was exposed to 30 minutes hypo-osmotic solution, followed by an additional 30 minutes in isotonic solution. Top images (rainbow pseudocolor) are FLIM images of live cells. Bottom images (red) are lamin A/C immunostaining of fixed samples. **B)** Quantification of FRET efficiency for hypo-osmotic shock experiments. **C)** Quantification of nuclear area changes for hypo-osmotic shock experiments. **D)** HeLa cells with gp210 tension sensor knock-in were exposed to isotonic (control) or hyper-osmotic buffer to induce nuclear shrinking. Additionally, a recovery sample was exposed to 30 minutes hyper-osmotic solution, followed by an additional 30 minutes in isotonic solution. **E)** Quantification of FRET efficiency for hyper-osmotic shock experiments. **F)** Quantification of nuclear area changes for hyper-osmotic shock experiments.

When cells subjected to hypo-osmotic shock were returned to an isotonic solution (recovery), we observed an increase in FRET, indicating that NPC forces are reduced once nuclear swelling is stopped. However, immunostaining of lamin A/C showed that recovered nuclei had significantly reduced nuclear area (**Figure 2C**) along with altered nuclear morphology (**Figure 2A, Supplemental Figure 3**) as compared to unstimulated (control) cells, indicating that hypo-osmotic swelling is not a purely elastic deformation, but may have viscous or plastic components. We also observed a higher gp210 TS FRET efficiency following recovery as compared to the FRET efficiency observed prior to osmotic shock (**Figure 2C**), indicating that NPC forces are also reduced in recovered cells.

In parallel experiments we also tested the effects of hyper-osmotic shrinking experiments. Hyper-osmotic shock decreased nuclear area (**Figure 2D, 2F**). Additionally, hyper-osmotic shock increased gp210TS FRET efficiency, indicating a decrease in mechanical force across the NPC (**Figure 2E**). A more complete recovery for gp210TS was observed during nuclear shrinking, which suggests that nuclear shrinking may be a more elastic phenomena than nuclear swelling (**Figure 2F**).

Two control sensors were developed to further validate gp210 TSmod FRET-force changes: gp210-TS F7 and gp210Sec61β TSmod (**Supplemental Figure 4A**). gp210-TS F7 has the original 40 amino acid elastic linker in TSmod truncated to 7 amino acids (termed F7), as the shorter F7 linker has been reported to be unable to elongate under force.^11^ The gp210-Sec61β sensor had the transmembrane domain of gp210 replaced with Sec61β to inhibit the localization of the gp210 sensor to the NPC.^10^ gp210-Sec61β TSmod contains the full length (40 amino acid) elastic linker of TSmod, and may still report force changes across membranes outside of the NPC. Both control sensors showed similar localization to the nuclear envelope, as compared to the gp210 TSmod (**Supplemental Figure 4B**). As expected the F7 sensor showed higher FRET efficiency, due to the shorter linker (7 vs 40 amino acids) between the donor (mTFP1) and acceptor (venus) FRET pair. Unexpectedly, both the F7 and the gp210-Sec61β sensors exhibited similar changes with hypoosmotic shock (**Supplemental Figure 4C**). This result suggests that these control sensors are still responsive to the large mechanical deformations introduced by large osmotic stresses. We attempted to use more moderate levels of osmotic stress (1/3 water vs 2/3 water), but the gp210 TSmod did not respond to lower levels of osmotic stress (**Supplemental Figure 4C**).

### gp210 forces are sensitive to ECM stiffness

Given prior reports that NPC diameter is sensitive to ECM stiffness,^4^ we sought to understand how gp210 forces would differ during changes in cell spreading as well as changes in ECM stiffness. First, we compared the effects of cells spread on glass that was uncoated (control), coated with poly-L-lysine (PLL), or coated with fibronectin (FN). poly-L-lysine coated surfaces inhibit cell spreading and reduce the ability of focal adhesions to generate mechanical force.^7^ While there were no significant differences between cells grown on uncoated versus FN-coated surfaces, cells grown on PLL had significantly increased gp210TS FRET efficiency (**Figure 3A, 3B**), indicating that cell spreading and focal adhesion forces increases the mechanical tension on gp210. Additionally, the gp210-TS F7 and gp210/Sec61-β control sensors were not responsive to cell spreading, showing no significant difference in FRET efficiency on control or PLL-coated surfaces (**Supplemental Figure 4D**).

**Figure 3:**
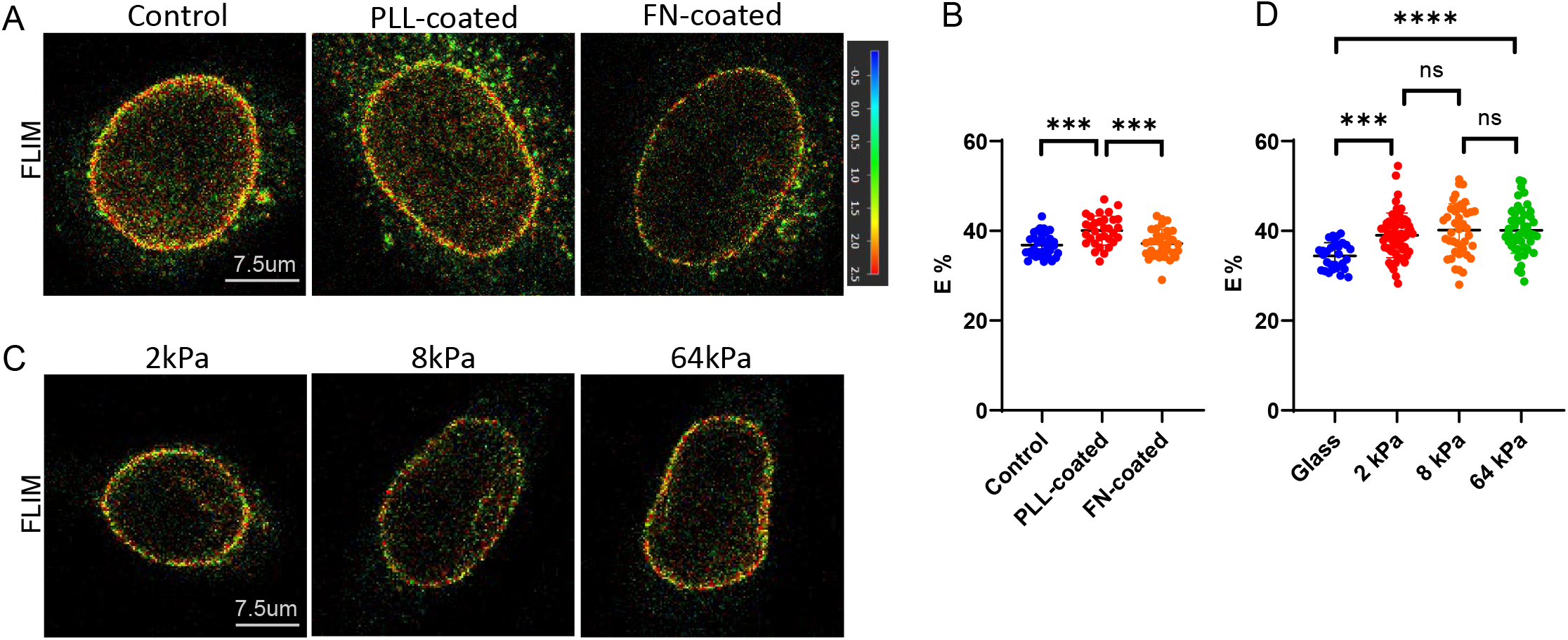
gp210 forces are sensitive to changes in ECM. **A)** HeLa cells with gp210 tension sensor knock-in were seeded on glass coverslips that were uncoated (control) or coated with poly-L-lysine (PLL) or fibronectin (FN). Images are FLIM images of live cells. **B)** FRET efficiency was highest for PLL coated surfaces, indicating that gp210 forces are lowest in unspread cells. No differences were observed between uncoated or FN-coated surfaces. **C)** Cells were seeded onto polyacrylamide gels with elastic moduli of 2kPa, 8kPa, or 64kPa and compared to cells seeded on glass. Images are FLIM images of live cells. **D)** While cells grown on glass had a reduced FRET efficiency compared to polyacrylamide gels, there were no differences between the polyacrylamide gels of varying stiffness.

Next, we compared gp210 FRET efficiency on cells cultured on fibronectin coated 2kPa, 8kPa, and 64 kPa polyacrylamide substrates. We did not observe any differences in gp210 FRET efficiency across these substrates (**Figure 3D, 3E**). When compared to cells cultured on glass (∼1000 MPa), we observed that gp210TS FRET efficiency was significantly less on glass when compared to the polyacrylamide substrates (**Figure 3D, 3E**).

### NPC forces are dependent on the nuclear LINC complex

Because the LINC complex was shown to be important for changes in the mechanosensitive transport of YAP,^4^ we investigated how disruption of the LINC complex affects gp210 forces. To disrupt the LINC complex we used siRNA designed to knockdown SUN1 and SUN2.^16^ This approach resulted in a large decrease in SUN1 and SUN2 protein expression (**Figure 4A**).

**Figure 4:**
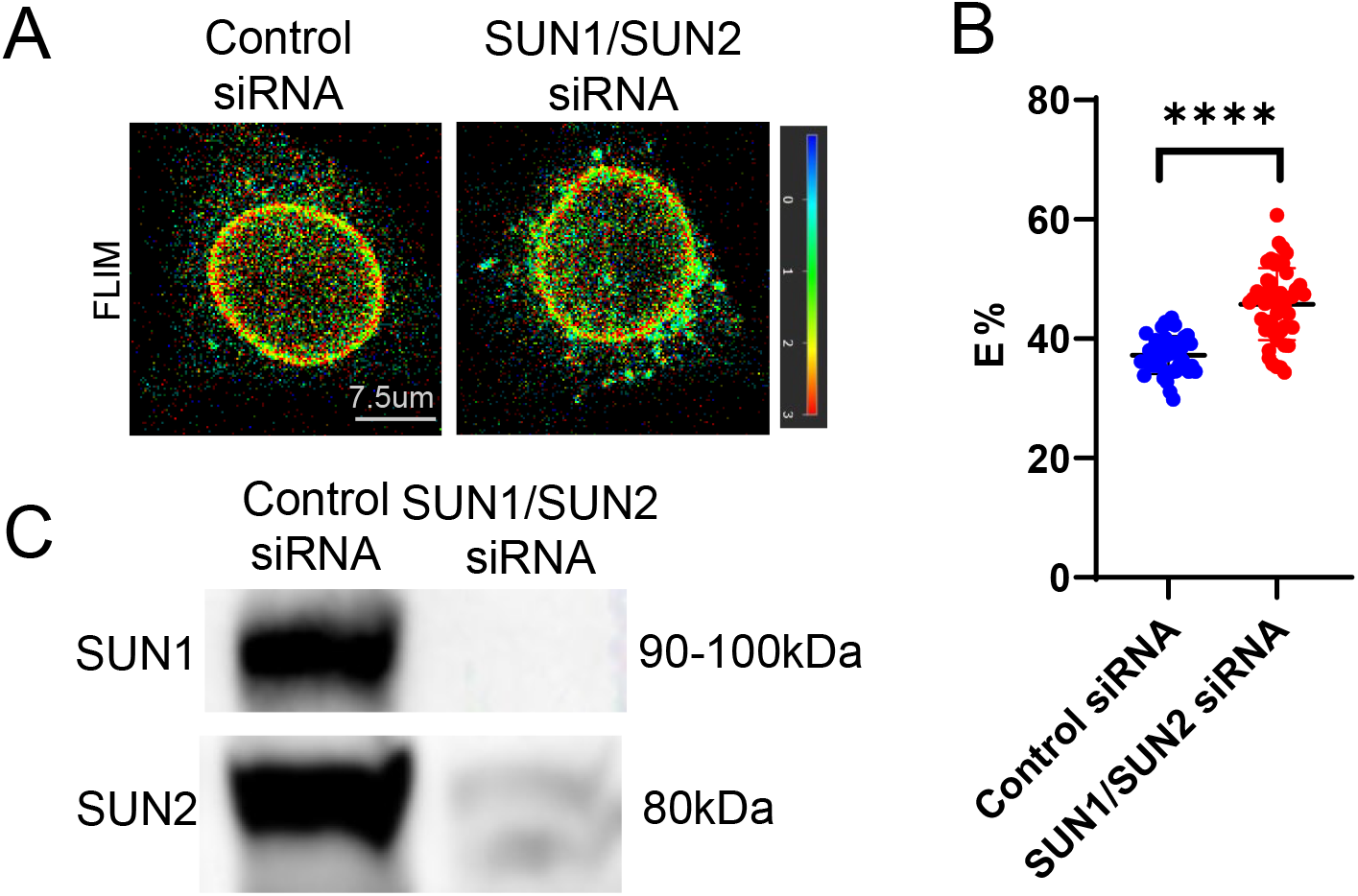
gp210 forces are dependent on the nuclear LINC complex. **A)** HeLa cells with the gp210 sensor knock-in were transfected with control siRNA (non-silencing) or siRNA targeted to SUN1 and SUN2. SUN1 and SUN2 siRNA treatment resulted in a large decrease in SUN1 and SUN2 protein expression. **B)** FLIM images of the gp210 tension sensor in control and SUN1/2 siRNA treated cells. **C)** Quantification of FRET efficiency showed that knockdown of SUN1 and SUN2 results in increased FRET efficiency for the gp210 sensor.

Next, we examined how depletion of SUN1 and SUN2 affected gp210 TS FRET (**Figure 4B**). Quantification of FRET showed that loss of SUN1 and SUN2 increased gp210 FRET (**Figure 4C**), indicating that disruption of the LINC complex reduces gp210 forces in resting cells.

### NPC forces are dependent on chromatin condensation

Changes in chromatin compaction have been shown to affect overall nuclear shape, volume, and stiffness.^18–20^ We therefore used Tricostatin A (TSA) and methylstat to decondense or condense (respectively) chromatin (**Figure 5A**). Decondensation of chromatin with TSA decreased gp210 FRET, indicating increased tension on gp210 (**Figure 5B**). However, condensation of chromatin with methylstat did not significantly affect FRET, indicating further compaction of chromatin did not affect gp210 force (**Figure 5B**). The gp210-TS F7 control sensor did not respond to TSA treatment, further confirming the force-responsiveness of the gp210 TSmod (**Supplemental Figure 4E**). Interestingly the gp210/Sec61-β control sensor exhibited an opposite change in FRET (increased FRET efficiency) (**Supplemental Figure 4E**).

**Figure 5:**
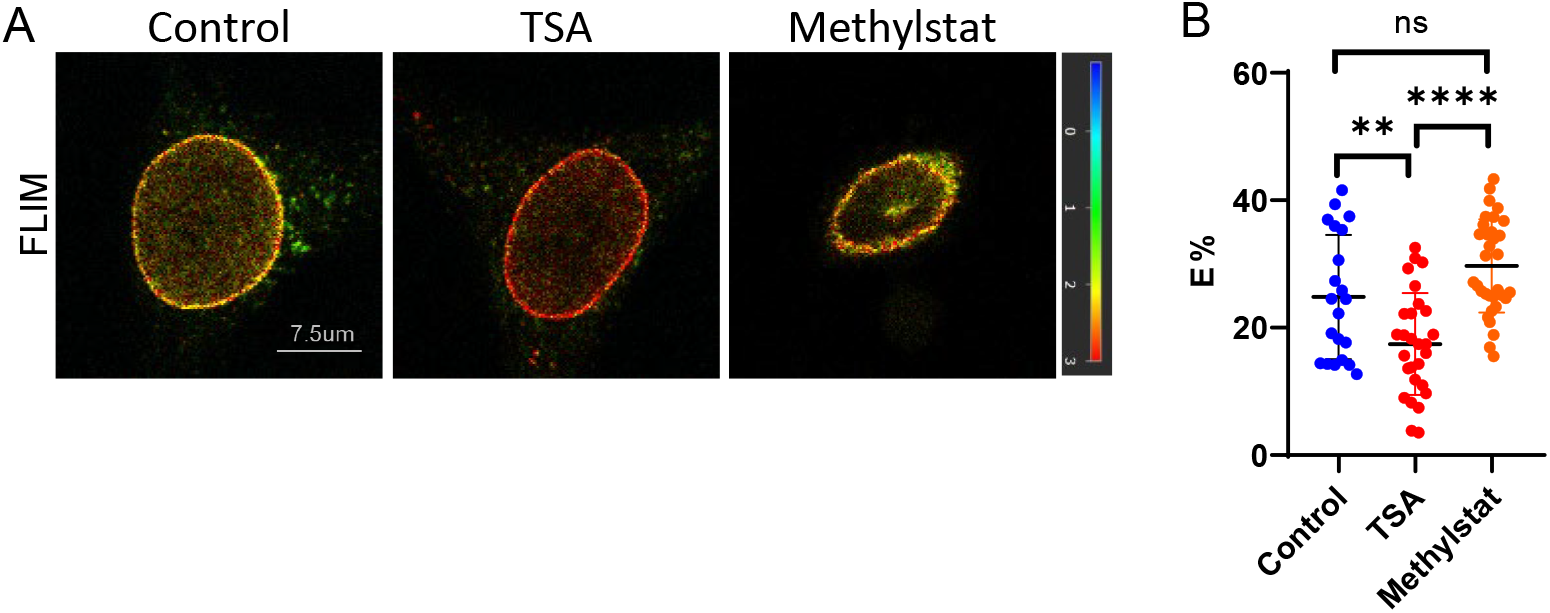
gp210 forces are dependent on chromatin condensation. **A**) FLIM images of HeLa cells with the knock-in gp210 sensor that were untreated (control) or treated with Tricostatin A (TSA) or methylstat. **B)** TSA resulted in a significant decrease in gp210 sensor FRET efficiency whereas methylstat resulted in no significant difference in FRET efficiency when compared to control cells.

### gp210 forces are sensitive to actomyosin contractility

Because forces on nuclear LINC complex proteins have been shown to be influenced by actomyosin contractility,^8,9^ we sought to understand how perturbations in actomyosin contractility would influence gp210 force (**Figure 6A**). Unexpectedly, inhibition of MLCK (using ML7) resulted in a decrease in gp210 sensor FRET, indicating increased force (**Figure 6B**).However, inhibition of ROCK (using Y-27632) did not affect gp210 FRET (**Figure 6B**). Through careful measurements of nuclear elongation, we observed that ML7, but not Y-27632, resulted in increased nuclear elongation (**Figure 6C**). No increase in nuclear height was observed for ML7 treatment (**Figure 6D**). Thus, it may be possible that ML7-induced nuclear morphology changes are responsible for the increased gp210 forces. The differences between ML7 and Y-compound with regard to nuclear morphology are supported by a previous study which also observed stronger effects of ML7 on nuclear flattening as compared to Y-27632.^21^ Activation of RhoA, using Rho Activator II, resulted in increased gp210 sensor FRET, indicating reduced force on gp210 (**Figure 6B**). Rho Activator II also resulted in increased nuclear elongation (**Figure 6C**). The reasons for the reduced gp210 force in response to increased active RhoA are unclear. The gp210-TS F7 and gp210/Sec61-β control sensors were not responsive to ML7 (**Supplemental Figure 4F**).

**Figure 6:**
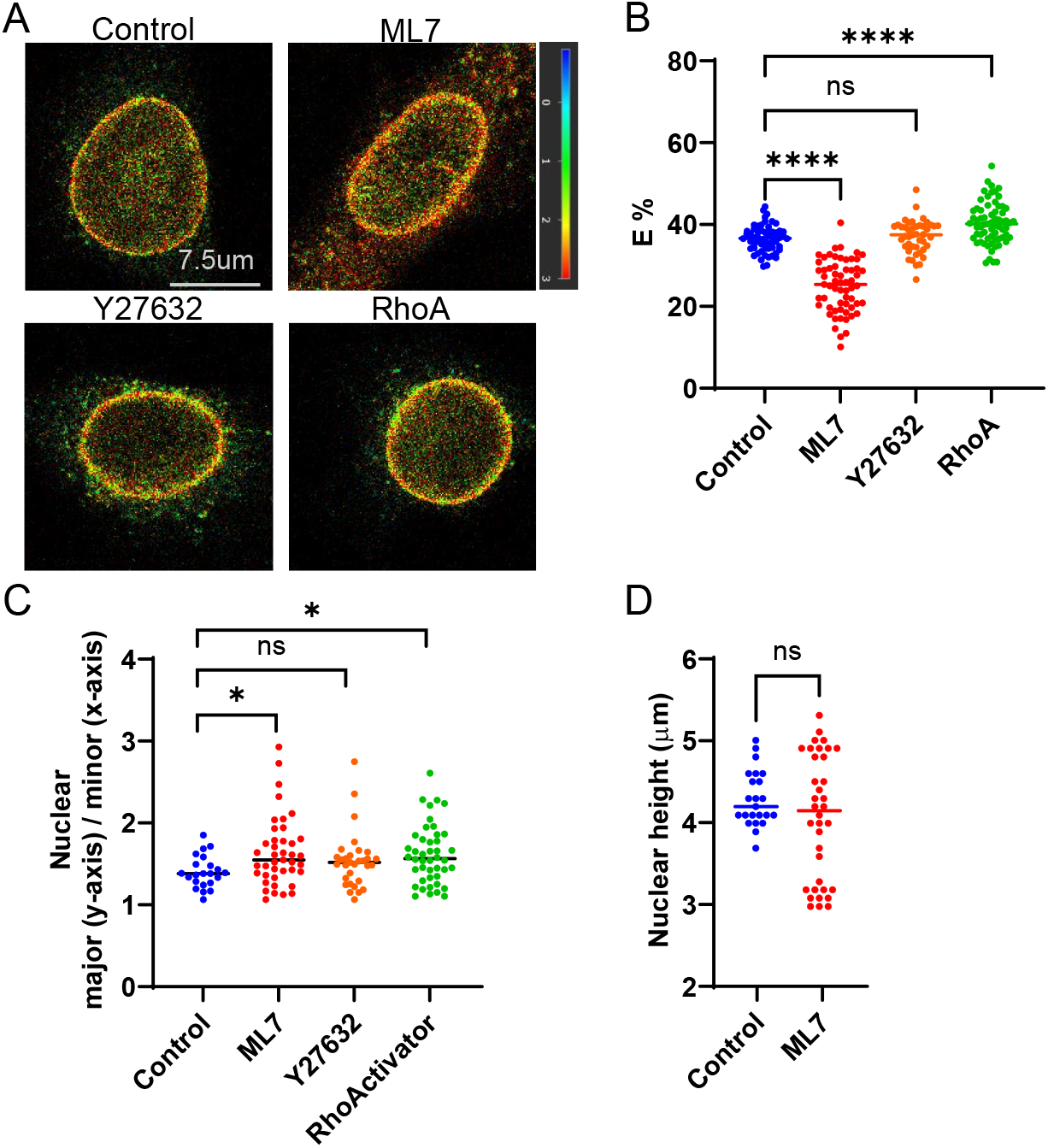
gp210 forces are influenced by myosin contractility. **A)** FLIM images of HeLa cells with the knock-in gp210 sensor that were untreated (control) or treated with ML7, Y-27632, or RhoA activator II. **B)** Quantification of FRET efficiency for each condition showed that ML7 but not Y27632 significantly reduced gp210 sensor FRET efficiency. In contrast, Rho Activator II increased FRET efficiency for the gp210 sensor. **C)** Analysis of nuclear height for all conditions showed that ML7 and Rho Activator II significantly increased nuclear elongation, whereas Y27632 did not affect nuclear elongation. **D)** Nuclear height was not affected by ML7 treatment.

Recent work has shown that HeLa cells have NPC with larger diameters at the bottom of the nucleus, as compared to the top surface.^22^ This finding, coupled with prior studies, ^23,24^ have suggested that the bottom of the nucleus may experience higher levels of strain. Thus, we sought to understand if there are spatial differences in gp210 force across the surface of the nucleus. Initially we sought to analyze gp210 FRET at the top, middle, and bottom surfaces of individual nuclei. However, the high curvature of the top of the nucleus resulted in images which had fewer fluorescent pixels (reducing FLIM counts), resulting in large variations in the calculated lifetime. Therefore, we only compared gp210 FRET at the middle (lateral/equatorial) regions of the nucleus to the bottom (basal) surface. We did not observe any significant differences in FRET efficiency between these two regions (**Supplemental Figure 5**). We are unable to determine if the lack of a difference is due to a lack of difference in mechanical tension on gp210 or a lack of sensitivity in our biosensor for detecting small changes in force.

### Increased gp210 tension is correlated with increased protein export from the nucleus

Prior work has shown that ECM stiffness drives increased import and export rates for proteins passing through the NPC.^5^ Thus we speculated that conditions which increase gp210 forces, such as hypoosmotic shock, would also affect nuclear export rates. Using an existing optogenetic nuclear export sensor, known as LEXY,^15^ we measured nuclear export in cells in isotonic and hypotonic conditions (**Figure 7A**). The rate of export was higher under hypotonic conditions (**Figure 7B**), indicating that increases in gp210 forces are correlated to increased export.

**Figure 7.**
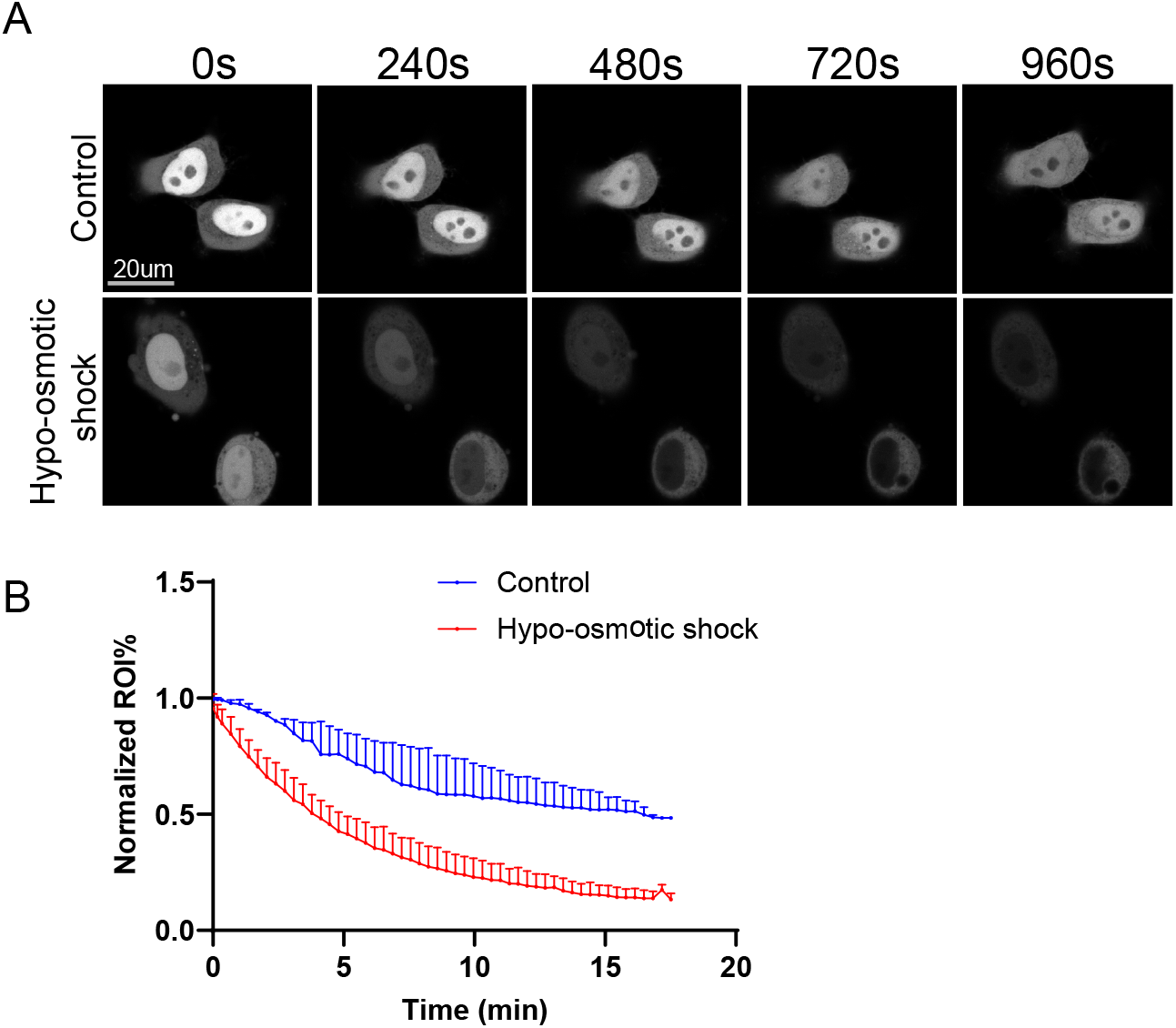
Hypoosmotic shock increases export of proteins from the nucleus. **A)** Using an existing LEXY optogenetic export sensor, the rate of nuclear export was measured in cells cultured in isotonic and hypotonic conditions. The sensor was photoactivated for export with 488 nm light at time=0 seconds. Images were taken at the indicated time intervals. **B)** Quantification of nuclear LEXY signal over time. Nuclear fluorescent intensity was normalized to the starting value prior to light activation.

## Discussion

Although the NPC has been hypothesized to experience mechanical force, to date there have not yet been any direct measurements of protein-level forces on the NPC. In this work we developed a gp210 FRET-force sensor, demonstrating that this transmembrane nucleoporin experiences significant mechanical tension. Our work suggests that nuclear strain is a significant regulator of gp210 tensile forces. We note that the increased forces reported by the gp210 tension sensor are correlated to functional changes in the NPC (increased nuclear-cytosolic transport, **Figure 7**). Future studies will be needed to better understand how NPC forces change during other biological processes, including if there is spatial variation in NPC forces.

To validate the FRET-force responsiveness of our new gp210 force sensor we developed two control sensors: gp210-TS F7 (which has a shorter rigid peptide between the FRET pair) and gp210/Sec61-β (which is not localized to NPCs). While gp210 TS showed decreased FRET (increased force) during cell spreading (**Figure 3B**) and ML-7 treatment (**Figure 6B**), both control sensors were insensitive to cell spreading (**Figure 3C**) and ML-7 treatment (**Figure 6E**). Additionally, while chromatin relaxation with TSA also decreased gp210 FRET (**Figure 5B**), the gp210 F7 control sensor did not show any change (**Figure 5C**). However, the gp210/Sec61-β showed increase FRET in response to TSA. Because the gp210/Sec61-β control sensor is localized outside of the NPC, the FRET changes observed may be a result of changes in forces across the nuclear envelope and surrounding ER. Notably, the opposite responses of gp210TS (decreased FRET, increased force) and gp210/Sec61-β (increased FRET, decreased force) to TSA treatment provide strong evidence that the gp210 sensor can distinguish NPC-specific forces from general nuclear envelope/ER mechanical changes. This dissociation is consistent with a model in which chromatin decompaction increases nuclear volume and nuclear envelope strain, thereby increasing tension on pore-localized gp210, while simultaneously altering the mechanical environment of non-pore nuclear envelope/ER proteins differently. Finally, these two control sensors exhibited similar FRET decreases in response to hypoosmotic stress. It is not clear why osmotic stress reduces the FRET in these control sensors. In the case of the gp210/Sec61-β control sensor, it may be possible that hypoosmotic stress is imparting tension across the nuclear envelope and surrounding ER. We did observe a significant decrease in FRET for both the nuclear envelope and ER regions of this sensor (**Supplemental Figure 4C**), suggesting that the ER also experiences mechanical stretching during osmotic shock. In the case of the gp210 F7 control sensor, the decrease in FRET may occur from large mechanical forces driving unfolding of the fluorescent proteins in TSmod.^25,26^ In summary the gp210-Sec61β control sensor can be used to identify when forces are NPC-specific, and the gp210 F7 sensor serves as a control to understand if treatment conditions may be unfolding or otherwise altering the gp210 sensor itself. An important consideration is that the gp210TS knock-in sensor is expressed at endogenous levels, whereas the F7 and gp210-Sec61β control sensors are transiently overexpressed. While differences in expression level could affect absolute FRET baselines, the lack of responsiveness of both control sensors to ML7 and cell spreading perturbations supports the conclusions that FRET changes observed with the endogenous gp210TS reflect force-specific responses.

Our results suggest that strain on the nuclear envelope may be an important driver of gp210 forces, in addition to actomyosin-based forces applied through the LINC complex. For example, decondensation of chromatin (**Figure 5**) resulted in increased gp210 forces. This contrasts with prior work where we showed that chromatin decondensation reduced lamin A/C forces.^9^ Similarly, inhibition of myosin light chain kinase resulted in increased gp210 forces (**Figure 6**), which also contrasts with our prior work where we showed that actomyosin contractility inhibition reduces nesprin-2G^8^ and lamin A/C forces.^9^ These results indicate that NPC forces may be dependent on both forces applied through the LINC complex, as well as nuclear morphology changes that alter the strain on the nuclear envelope.

Interestingly, gp210 is not expressed in all cell types, suggesting that gp210 has cell-type specific functions.^27^ Thus an intriguing possibility is that expression of gp210 alters the mechanical forces on the NPC. Supporting this hypothesis, a prior study showed that antibodies that bind the lumenal domain of gp210 can inhibit NPC dilation.^28^ Future studies will be needed to explore how expression of gp210 affects NPC transport kinetics. We also report that we attempted to develop TSmod sensors for ndc1 and pom151, the other two transmembrane nucleporins which are more ubiquitously expressed, by inserting Tsmod at various positions in these proteins. However, these sensors did not localize properly to the nuclear envelope (not shown).

One limitation of our nuclear export experiments (**Figure 7**) is the use of hypo-osmotic shock, which our control sensor data indicates changes FRET through mechanisms beyond NPC-specific tension. Additionally, hypo-osmotic swelling increases nuclear volume, which could independently affect apparent export kinetics by diluting nuclear contents. Thus, while our data show a correlation between conditions that increase gp210 forces and increased export rates, future studies using perturbations where the control sensors are unresponsive (such as ML7 or TSA) will be needed to more directly link NPC-specific mechanical tension to changes in transport kinetics.

In summary this work demonstrates that gp210, a transmembrane nucleoporin, is subject to tensile loads. The gp210 sensor was able to resolve changes in force under various perturbations. We anticipate that the gp210 FRET-force sensor will be a useful tool for future studies of the mechanobiology of the NPC and nuclear-cytoskeletal transport.

## Acknowledgements

We acknowledge support from Dario Palmieri and Lara Rizzotto in the OSU Gene Editing Shared Resource for assisting with designing and developing the CRISPR strategy for the gp210 knock-in cell line. This shared resource is supported in part by funding from the National Institute of Health award P30 CA016058.

We acknowledge helpful conversations with Teemu Ihalainen throughout this project.

The studies were supported by funding from the National Institute of Health award R35 GM119617 (to D.E.C.) and National Science Foundation award CMMI 2234888 (to D.E.C.) as well as the Paul G. Allen Frontiers Group of the Paul G. Allen Family Foundation, and Allen Distinguished Investigator Award (to G.W.G.L.).

## Figure Legends

**Supplemental Figure 1. Acceptor photobleaching of transiently transfected gp210 TSmod**. HeLa cells were transiently transfected with rat gp210 TSmod. Cells were subjected to acceptor photobleaching. Increased donor fluorescence was observed following acceptor photobleaching, confirming FRET.

**Supplemental Figure 2. FLIM analysis of transiently transfected gp210 TSmod. A)** HeLa cells were transiently transfected with rat gp210 TSmod and subjected to isotonic (control) or hypo-osmotic conditions. Additionally, cells exposed to 30 minutes of hypo-osmotic media were changed back to isotonic media for approximately 30 minutes. Top (blue) images are fluorescence intensity, bottom row are FLIM images. **B)** Quantification of FRET efficiency for hypo-osmotic shock experiments.

**Supplemental Figure 3. Immunostaining of cells during hypo-osomotic shock and recovery A)** Cells were fixed and immunostained with anti-NPC and anti-lamin A antibodies for the indicated conditions and timepoints. Nuclei and actin did not fully recover after 15 minutes of recovery in isotonic media. **B)** Cells were fixed and immunostained with anti-vimentin and anti-lamin A antibodies. Vimentin did not fully recover after 15 minutes of recovery in isotonic media.

**Supplemental Figure 4. Design of gp210 control (force-insensitive) sensors. A)** Schematic of the gp210 TSmod force sensor (gp210-TS), F7, and the gp210 sensor with the Sec61β transmembrane domain. **B)** Fluorescent images of cells transfected with each sensor. All three sensors were localized to the nuclear envelope. **C)** gp210-TS, F7, and gp210-Sec61β all exhibited significant FRET increases in response to hypo-osmotic shock. **D)** FRET efficiency of gp210-TS, F7, and gp210-Sec61β sensors of spread cells or cells seeded on poly-L-lysine (to inhibit spreading). The F7 and gp210-Sec61β control sensors showed no differences between either condition. **E)** FRET efficiency of gp210-TS, F7, and gp210-Sec61β control sensors with or without TSA treatment. The gp210-TS F7 control sensor did not respond to TSA, while the gp210-Sec61β control sensor showed increased FRET efficiency with TSA.**F)** FRET efficiency of gp210-TS, F7, and gp210/Sec61β control sensors with or without ML7 treatment. Neither control sensor showed a significant change in FRET efficiency with ML7.

**Supplemental Figure 5. Analysis of gp210 FRET differences between the middle and bottom surfaces of the nucleus. A)** FLIM images from the middle and bottom (basal) surfaces of the same nucleus. **B)** FRET efficiency from paired images. No significant difference was observed between middle and the bottom surfaces of the nucleus (paired t-test).

**Supplemental Video 1:** Using an existing LEXY optogenetic export sensor, the rate of nuclear export was measured in cells cultured in isotonic conditions. The sensor was photoactivated for export with 488 nm light at time=0 seconds. Images were taken every 150 seconds (1 frame=150 s).

**Supplemental Video 2:** Using an existing LEXY optogenetic export sensor, the rate of nuclear export was measured in cells cultured in hypotonic conditions. The sensor was photoactivated for export with 488 nm light at time=0 seconds. Images were taken every 150 seconds (1 frame=150 s).

## References

1. Matsuda, A., and Mofrad, M.R.K. (2022). On the nuclear pore complex and its emerging role in cellular mechanotransduction. APL Bioeng. 6, 011504. 10.1063/5.0080480.

2. Wolf, C.B., and Mofrad, M.R.K. (2009). Mechanotransduction. In Cellular Mechanotransduction (Cambridge University Press), pp. 417–437. 10.1017/CBO9781139195874.019.

3. Wang, N., Tytell, J.D., and Ingber, D.E. (2009). Mechanotransduction at a distance: Mechanically coupling the extracellular matrix with the nucleus. Nat. Rev. Mol. Cell Biol. 10, 75–82. 10.1038/NRM2594,.

4. Elosegui-Artola, A., Andreu, I., Beedle, A.E.M., Lezamiz, A., Uroz, M., Kosmalska, A.J., Oria, R., Kechagia, J.Z., Rico-Lastres, P., Le Roux, A.-L., et al. (2017). Force Triggers YAP Nuclear Entry by Regulating Transport across Nuclear Pores. Cell 171, 1397-1410.e14. 10.1016/j.cell.2017.10.008.

5. Andreu, I., Granero-Moya, I., Chahare, N.R., Clein, K., Molina-Jordán, M., Beedle, A.E.M., Elosegui-Artola, A., Abenza, J.F., Rossetti, L., Trepat, X., et al. (2022). Mechanical force application to the nucleus regulates nucleocytoplasmic transport. Nat. Cell Biol. 24, 896–905. 10.1038/s41556-022-00927-7.

6. Zimmerli, C.E., Allegretti, M., Rantos, V., Goetz, S.K., Obarska-Kosinska, A., Zagoriy, I., Halavatyi, A., Hummer, G., Mahamid, J., Kosinski, J., et al. (2021). Nuclear pores dilate and constrict in cellulo. Science (1979). 374. 10.1126/SCIENCE.ABD9776,.

7. Grashoff, C., Hoffman, B.D., Brenner, M.D., Zhou, R., Parsons, M., Yang, M.T., McLean, M.A., Sligar, S.G., Chen, C.S., Ha, T., et al. (2010). Measuring mechanical tension across vinculin reveals regulation of focal adhesion dynamics. Nature 466, 263–266. 10.1038/nature09198.

8. Arsenovic, P.T., Ramachandran, I., Bathula, K., Zhu, R., Narang, J.D., Noll, N.A., Lemmon, C.A., Gundersen, G.G., and Conway, D.E. (2016). Nesprin-2G, a Component of the Nuclear LINC Complex, Is Subject to Myosin-Dependent Tension. Biophys. J. 110, 34–43. 10.1016/j.bpj.2015.11.014.

9. Danielsson, B.E., George Abraham, B., Mäntylä, E., Cabe, J.I., Mayer, C.R., Rekonen, A., Ek, F., Conway, D.E., and Ihalainen, T.O. (2023). Nuclear lamina strain states revealed by intermolecular force biosensor. Nat. Commun. 14, 3867. 10.1038/s41467-023-39563-6.

10. Gomez-Cavazos, J.S., and Hetzer, M.W. (2015). The nucleoporin gp210/Nup210 controls muscle differentiation by regulating nuclear envelope/ER homeostasis. J. Cell Biol. 208, 671. 10.1083/JCB.201410047.

11. Kanoldt, V., Kluger, C., Barz, C., Schweizer, A.-L., Ramanujam, D., Windgasse, L., Engelhardt, S., Chrostek-Grashoff, A., and Grashoff, C. (2020). Metavinculin modulates force transduction in cell adhesion sites. Nat. Commun. 11, 6403. 10.1038/s41467-020-20125-z.

12. Mills, J., Tessari, A., Anastas, V., Kumar, D.S., Rad, N.S., Lamba, S., Cosentini, I., Reers, A., Zhu, Z., Miles, W.O., et al. (2024). Nucleolin acute degradation reveals novel functions in cell cycle progression and division in TNBC. bioRxiv. 10.1101/2024.06.17.599429,.

13. Capece, M., Tessari, A., Mills, J., Vinciguerra, G.L.R., Louke, D., Lin, C., McElwain, B.K., Miles, W.O., Coppola, V., Davies, A.E., et al. (2023). A novel auxin-inducible degron system for rapid, cell cycle-specific targeted proteolysis. Cell Death Differ. 30, 2078–2091. 10.1038/S41418-023-01191-4,.

14. Gutschner, T., Haemmerle, M., Genovese, G., Draetta, G.F., and Chin, L. (2016). Post-translational Regulation of Cas9 during G1 Enhances Homology-Directed Repair. Cell Rep. 14, 1555–1566. 10.1016/j.celrep.2016.01.019.

15. Niopek, D., Wehler, P., Roensch, J., Eils, R., and Di Ventura, B. (2016). Optogenetic control of nuclear protein export. Nat. Commun. 7, 10624. 10.1038/ncomms10624.

16. Turgay, Y., Champion, L., Balazs, C., Held, M., Toso, A., Gerlich, D.W., Meraldi, P., and Kutay, U. (2014). SUN proteins facilitate the removal of membranes from chromatin during nuclear envelope breakdown. J. Cell Biol. 204, 1099–1109. 10.1083/JCB.201310116.

17. Rabut, G., Doye, V., and Ellenberg, J. (2004). Mapping the dynamic organization of the nuclear pore complex inside single living cells. Nat. Cell Biol. 6, 1114–1121. 10.1038/ncb1184.

18. Furusawa, T., Rochman, M., Taher, L., Dimitriadis, E.K., Nagashima, K., Anderson, S., and Bustin, M. (2015). Chromatin decompaction by the nucleosomal binding protein HMGN5 impairs nuclear sturdiness. Nat. Commun. 6. 10.1038/ncomms7138.

19. Stephens, A.D., Liu, P.Z., Banigan, E.J., Almassalha, L.M., Backman, V., Adam, S.A., Goldman, R.D., and Marko, J.F. (2018). Chromatin histone modifications and rigidity affect nuclear morphology independent of lamins. Mol. Biol. Cell 29, 220–233. 10.1091/mbc.E17-06-0410.

20. Damodaran, K., Venkatachalapathy, S., Alisafaei, F., Radhakrishnan, A. V., Jokhun, D.S., Shenoy, V.B., and Shivashankar, G. V. (2018). Compressive force induces reversible chromatin condensation and cell geometry–dependent transcriptional response. Mol. Biol. Cell 29, 3039–3051. 10.1091/mbc.E18-04-0256.

21. Li, Y., Lovett, D., Zhang, Q., Neelam, S., Kuchibhotla, R.A., Zhu, R., Gundersen, G.G., Lele, T.P., and Dickinson, R.B. (2015). Moving Cell Boundaries Drive Nuclear Shaping during Cell Spreading. Biophys. J. 109, 670. 10.1016/J.BPJ.2015.07.006.

22. Morgan, K.J., Carley, E., Coyne, A.N., Rothstein, J.D., Lusk, C.P., and King, M.C. (2025). Visualizing nuclear pore complex plasticity with pan-expansion microscopy. J. Cell Biol. 224. 10.1083/JCB.202409120/278050.

23. Carley, E., Stewart, R.K., Zieman, A., Jalilian, I., King, D.E., Zubek, A., Lin, S., Horsley, V., and King, M.C. (2021). The linc complex transmits integrin-dependent tension to the nuclear lamina and represses epidermal differentiation. Elife 10. 10.7554/ELIFE.58541.

24. Ihalainen, T.O., Aires, L., Herzog, F.A., Schwartlander, R., Moeller, J., and Vogel, V. (2015). Differential basal-to-apical accessibility of lamin A/C epitopes in the nuclear lamina regulated by changes in cytoskeletal tension. Nat. Mater. 14, 1252–1261. 10.1038/nmat4389.

25. Ham, T.R., Collins, K.L., and Hoffman, B.D. (2019). Molecular Tension Sensors: Moving Beyond Force. Curr. Opin. Biomed. Eng. 10.1016/j.cobme.2019.10.003.

26. Ganim, Z., and Rief, M. (2017). Mechanically switching single-molecule fluorescence of GFP by unfolding and refolding. Proc. Natl. Acad. Sci. U. S. A. 114, 11052–11056. 10.1073/PNAS.1704937114/SUPPL_FILE/PNAS.201704937SI.PDF.

27. Olsson, M., Schéele, S., and Ekblom, P. (2004). Limited expression of nuclear pore membrane glycoprotein 210 in cell lines and tissues suggests cell-type specific nuclear pores in metazoans. Exp. Cell Res. 292, 359–370. 10.1016/j.yexcr.2003.09.014.

28. Drummond, S.P., and Wilson, K.L. (2002). Interference with the cytoplasmic tail of gp210 disrupts “close apposition” of nuclear membranes and blocks nuclear pore dilation. J. Cell Biol. 158, 53–62. 10.1083/jcb.200108145.

