## Supplemental Figures 1-5 for "A novel FRET-force biosensor for nucleoporin gp210 reveals that the nuclear pore complex experiences mechanical tension"

- **Supplemental Figure 1. Acceptor photobleaching of transiently transfected gp210 TSmod.** HeLa cells were transiently transfected with rat gp210 TSmod. Cells were subjected to acceptor photobleaching. Increased donor fluorescence was observed following acceptor photobleaching, confirming FRET.
- **Supplemental Figure 2. FLIM analysis of transiently transfected gp210 TSmod. A)** HeLa cells were transiently transfected with rat gp210 TSmod and subjected to isotonic (control) or hypo-osmotic conditions. Additionally, cells exposed to 30 minutes of hypo-osmotic media were changed back to isotonic media for approximately 30 minutes. Top (blue) images are fluorescence intensity, bottom row are FLIM images. **B)** Quantification of FRET efficiency for hypo-osmotic shock experiments.
- **Supplemental Figure 3. Immunostaining of cells during hypo-osomotic shock and recovery A)** Cells were fixed and immunostained with anti-NPC and anti-lamin A antibodies for the indicated conditions and timepoints. Nuclei and actin did not fully recover after 15 minutes of recovery in isotonic media. **B)** Cells were fixed and immunostained with anti-vimentin and anti-lamin A antibodies. Vimentin did not fully recover after 15 minutes of recovery in isotonic media.
- **Supplemental Figure 4. Design of gp210 control (force-insensitive) sensors. A)** Schematic of the gp210 TSmod force sensor (gp210-TS), F7, and the gp210 sensor with the Sec61β transmembrane domain. **B)** Fluorescent images of cells transfected with each sensor. All three sensors were localized to the nuclear envelope. **C)** gp210-TS, F7, and gp210-Sec61β all exhibited significant FRET increases in response to hypo-osmotic shock. **D)** FRET efficiency of gp210-TS, F7, and gp210-Sec61β sensors of spread cells or cells seeded on poly-L-lysine (to inhibit spreading). The F7 and gp210-Sec61β control sensors showed no differences between either condition. **E)** FRET efficiency of gp210-TS, F7, and gp210-Sec61β control sensors with or without TSA treatment. The gp210-TS F7 control sensor did not respond to TSA, while the gp210-Sec61β control sensor showed increased FRET efficiency with TSA. **F)** FRET efficiency of gp210-TS, F7, and gp210/Sec61β control sensors with or without ML7 treatment. Neither control sensor showed a significant change in FRET efficiency with ML7.
- **Supplemental Figure 5. Analysis of gp210 FRET differences between the middle and bottom surfaces of the nucleus. A)** FLIM images from the middle and bottom (basal) surfaces of the same nucleus. **B)** FRET efficiency from paired images. No significant difference was observed between middle and the bottom surfaces of the nucleus (paired t-test).
- **Supplemental Video 1:** Using an existing LEXY optogenetic export sensor, the rate of nuclear export was measured in cells cultured in isotonic conditions. The sensor was photoactivated for export with 488 nm light at time=0 seconds. Images were taken every 150 seconds (1 frame=150 s).
- **Supplemental Video 2:** Using an existing LEXY optogenetic export sensor, the rate of nuclear export was measured in cells cultured in hypotonic conditions. The sensor was photoactivated for export with 488 nm light at time=0 seconds. Images were taken every 150 seconds (1 frame=150 s).

Supplemental Figure 1

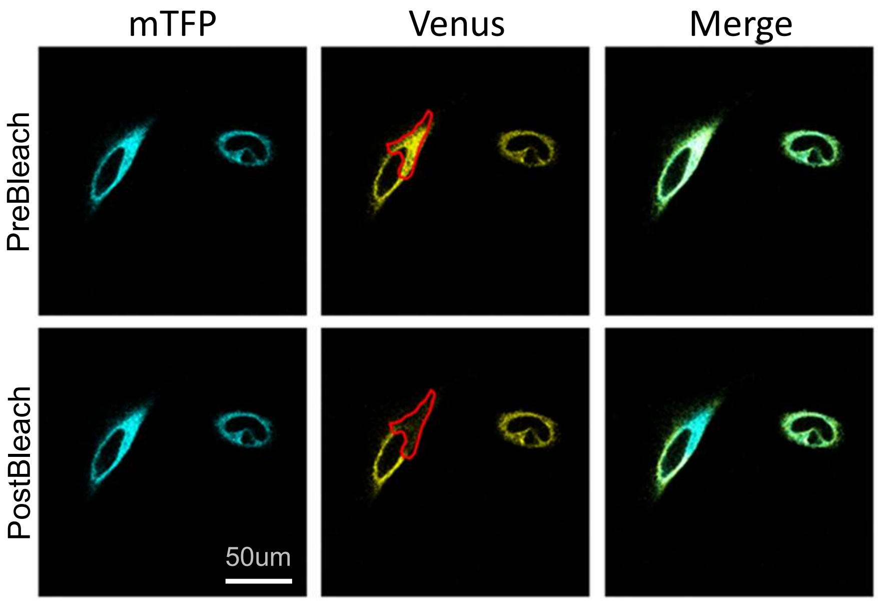

Supplemental Figure 2

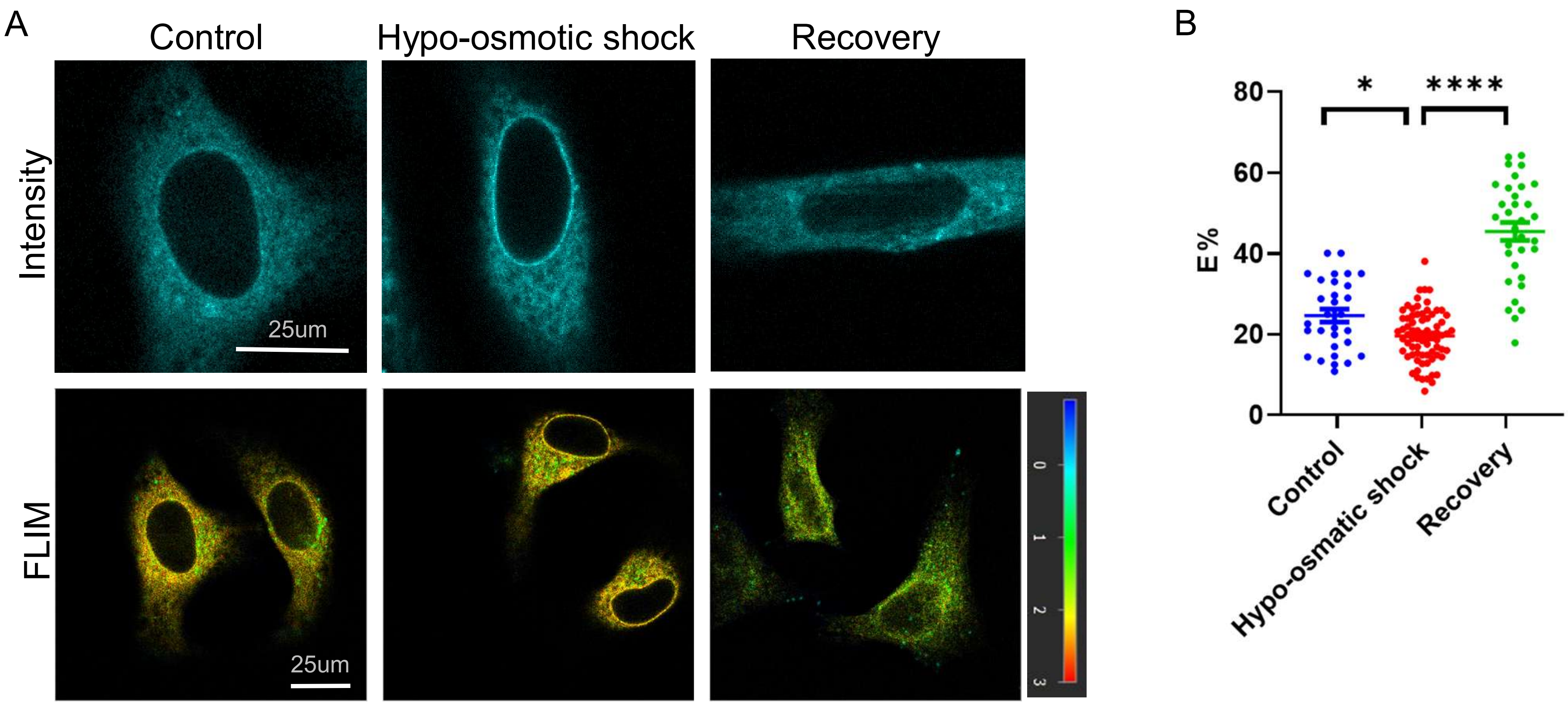

Supplemental Figure 3

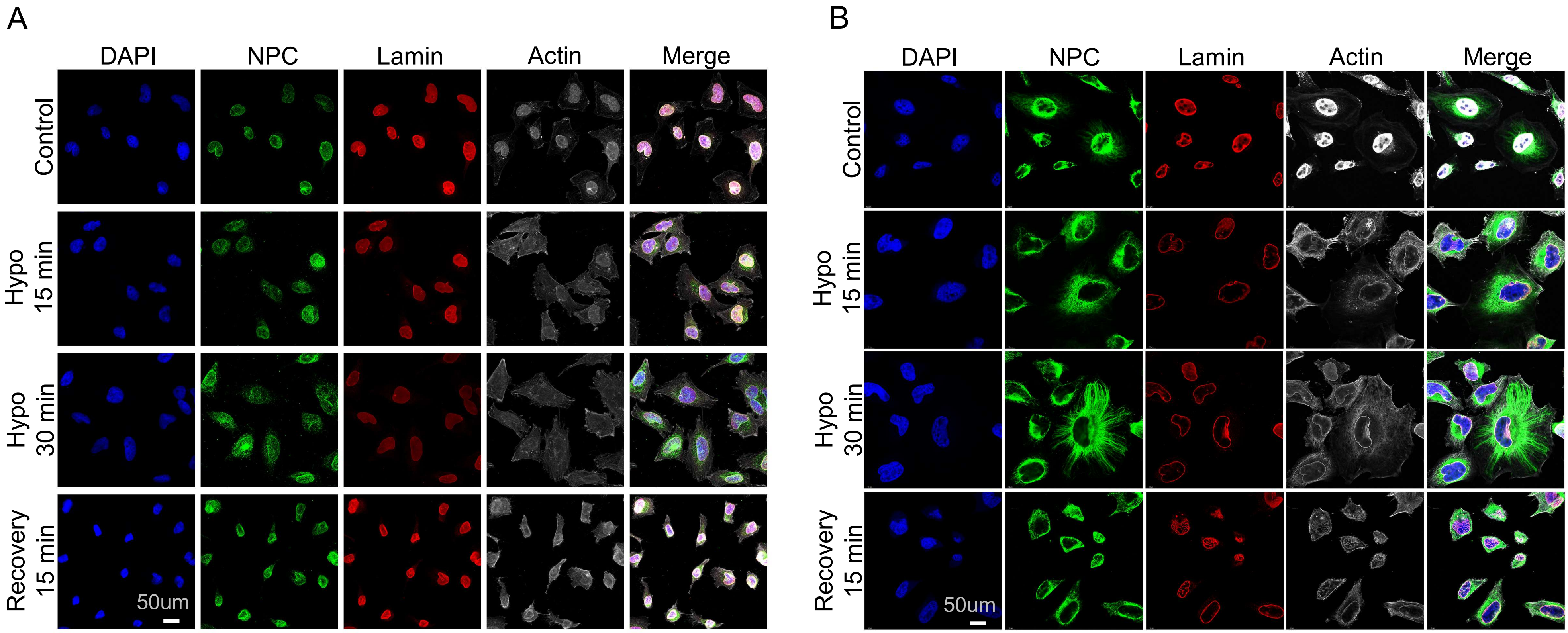

### Supplemental Figure 4

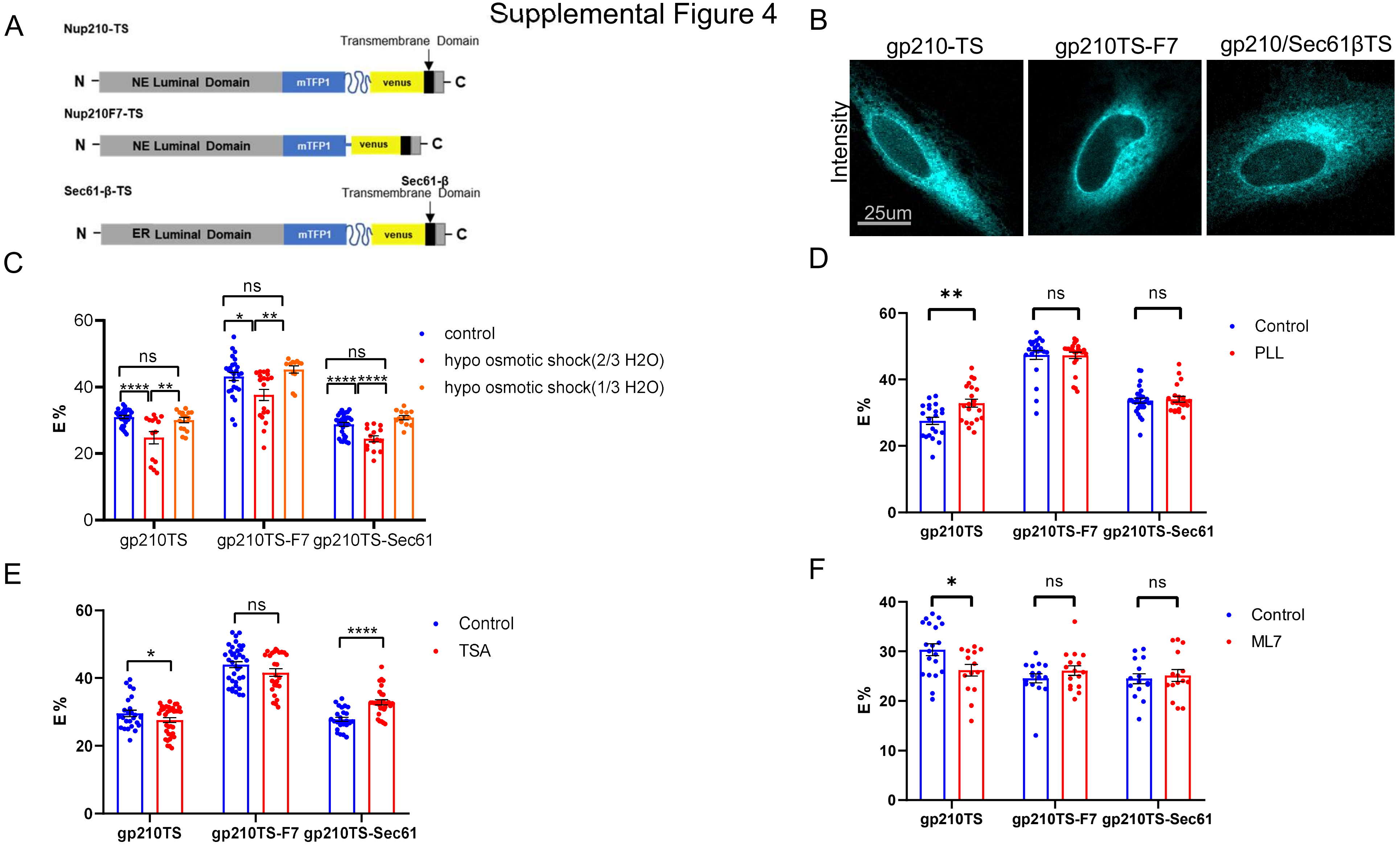

Supplemental Figure 5

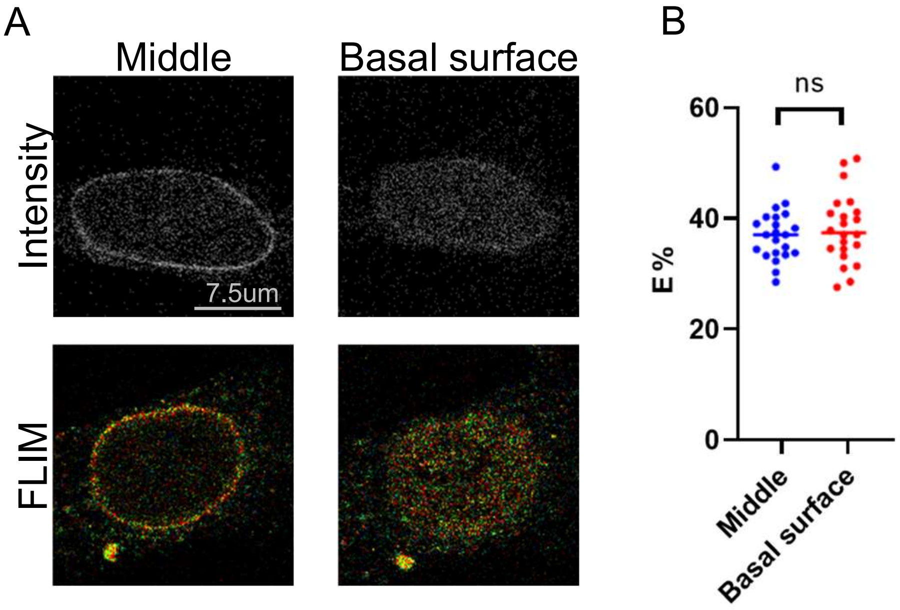
